# Early life gut microbiome dynamics mediate maternal effects on infant growth in vervet monkeys

**DOI:** 10.1101/2021.05.11.443657

**Authors:** Lauren Petrullo, Alice Baniel, Matthew J. Jorgensen, Sierra Sams, Noah Snyder-Mackler, Amy Lu

**Author notes:** Correspondence: Lauren Petrullo, Amy Lu.

## Abstract

**Background:** Maternal parity is associated with variation in infant growth across mammals, but the mechanisms underlying this relationship are unclear. Given emerging links between growth and the microbiome, and the importance of maternal microbiota in establishing this community, the assembly of the infant gut microbiome may be a mediator of parity effects on infant growth.

**Results:** Here, we analyzed 118 fecal and milk samples from mother-infant vervet monkey dyads across the first 6 months postpartum in a population with high growth-associated infant mortality. Despite poorer milk production, infants born to low parity females were larger at 6 months of age than their counterparts and exhibited divergent patterns in gut microbiome assembly. Gut microbiome alpha diversity increased rapidly from the first days of life to 4 months old in all infants, but infants born to low parity females exhibited reduced gut microbiome alpha diversity during early life. At the taxonomic level, infants broadly exhibited a shift from *Bacteroides fragilis* to *Prevotella* dominance. Infants of low parity females housed more *B. fragilis* in their guts, and *B. fragilis* dominance drove reduced alpha diversity. Maternal vertical transmission to the infant gut was greater from milk than from the maternal gut, and was greatest among infants born to low parity females. *B. fragilis* was 15-fold more abundant in milk than in the maternal gut and was greater in the milk of low parity females, suggesting that milk may be the primary maternal reservoir of *B. fragilis*. Path analyses demonstrated that both infant gut alpha diversity and *B. fragilis* mediated parity effects on postnatal growth: infants were larger at 6 months old if they exhibited reduced alpha diversity and a greater relative abundance of *B. fragilis* during early life.

**Conclusion:** The first days of life are a critical period of infant gut microbiome organization during which the establishment of a less diverse, milk-oriented microbial community abundant in *B. fragilis* promotes growth among infants born to reproductively inexperienced females.

## BACKGROUND

Across mammals, the pace of early infant growth has profound impacts on fitness. Stunted growth can result in delayed maturation, failure to recruit into the breeding population, and later-life disease [1–4]. Variation in postnatal growth is consistently associated with maternal traits in a number of species, suggesting that the maternal environment is a key factor influencing early growth. For example, infants born to reproductively inexperienced mothers typically grow slower, are smaller in body mass for their age, and exhibit higher rates of mortality than infants born to multiparous females [5–10]. Low parity females also produce less milk than high parity females, with milk production increasing with each successive birth [11–17]. Given that mammalian infants rely entirely on milk to fuel somatic growth prior to weaning, the growth costs of maternal reproductive inexperience may be related to poor milk production. In some populations, however, infants of reproductively inexperienced females grow at similar rates as their counterparts despite lower milk volumes [18,19], suggesting that milk production alone may not fully explain the effects of maternal parity on infant growth. Emerging research suggests that the early organization of the infant gut microbiome may mediate the link between parity and early postnatal growth. In humans and rodent models, reduced alpha diversity has been associated with growth deficits [20]; however, the effects have not been consistent, with other studies finding no relationship between microbial community diversity and growth [21]. In other animals, such as birds, the effects appear more consistent and with an opposite pattern: poultry birds with experimentally-reduced alpha diversity grew faster [22], and lower diversity during early life predicts faster later-life growth among free-living birds [23,24]. Beyond community diversity, experimental rodent models of the human microbiome have identified specific taxa are causally related to infant growth. For example, specialized microbes such as *Bacteroides fragilis* promote infant growth by enhancing host utilization of indigestible milk components such as glycans [21,25,26].

Importantly, the infant gut microbiome is organized through the vertical transmission of microbiota from maternal reservoirs (e.g., milk, fecal, vaginal, skin) [27], offering a potential avenue by which maternal traits such as parity may shape variation in this community. Early colonization by microbiota from maternal reservoirs determines not only the initial assembly of the infant gut microbiome, but subsequent patterns of microbial maturation [27,28]. However, this process of vertical transmission is not entirely stochastic: data in humans suggests that infants selectively seed specific microbes from highly diverse pools of maternal microbiota [29]. Preferential colonization of maternal bacteria may initiate the competitive exclusion of less favorable taxa [30], promoting microbiome-mediated developmental trajectories that reflect maternal interests, infant interests, or both [31,32]. Thus far, parity effects on the maternal microbial communities have been demonstrated in pigs and cows [33,34]. In pigs, maternal parity also influenced infant gut microbiome composition, evidence that parity effects on infant development may occur through variation in maternal vertical transmission. However, whether growth-related microbiome measures such as infant alpha diversity and *B. fragilis* abundance vary with maternal parity is unknown.

Importantly, three major maternal reservoirs may be important in coordinating maternal effects on variation in infant gut microbiome composition: the maternal gut and vaginal microbiomes, which seed the infant gut around parturition, and the maternal milk microbiome, which seeds the infant’s gut postnatally. Although recent studies suggest that the maternal gut is the largest source of colonizing bacteria to the infant gut [28,35– 37], consideration of the milk microbiome as a source of bacteria to the infant gut remains absent from these studies. Similar to the lactational transfer of maternal hormones and immune factors, the transfer of bacteria from milk to the infant gut reflects a direct postnatal connection between maternal and infant physiologies. Milk harbors an ephemeral yet diverse community of microbiota [38–42] that contributes substantially to the colonization of the infant gut microbiome across postnatal life [43]. In humans, some microbial taxa are exclusively shared between milk and infant gut communities [44,45], evidence of a unique milk-infant gut transmission pathway independent of other maternal reservoirs. The transfer of microbiota from milk to the infant gut may therefore be a pathway by which maternal traits such as parity can influence infant developmental trajectories. Indeed, as mammary morphology varies with successive reproductions [46], the milk microbiome may be an additional component of mammary gland development that changes with parity.

Vervet monkeys *(Chlorocebus aethiops sabaeus)* are a valuable model system for investigating the relationship between parity, the infant gut microbiome, and early growth, particularly with respect to the role of the milk microbiome. Similar to humans, vervets possess a milk microbiome that is highly diverse and abundant in growth-associated taxa such as *Bacteroides fragilis* [41]. Variation in *Bacteroides* colonization in human infants is not explained by exposure to the vaginal microbiome [47], suggesting that the transfer of *B. fragilis* to infants may occur primarily via milk or the maternal gut. In this study, we use amplicon sequencing of the 16S rRNA gene to analyze matched milk and fecal microbiome samples mother-infant vervet monkey dyads in a captive population with a high rate of growth-associated infant mortality (>30% in the first month of life) [48,49]. This high mortality rate offers a unique opportunity to test our hypothesis in a system where infant growth is under strong selective pressure, but where subjects are also housed in a controlled environment.

Here, we test the hypothesis that the organization of the infant gut microbiome mediates the relationship between maternal parity and infant growth, and investigate whether maternal microbial communities contribute to this relationship. We predict that: 1) there is a direct effect of maternal parity on infant growth in this population; 2) maternal vertical transmission and the organization of the infant gut microbiome varies with parity; and 3) the effects of maternal parity on postnatal infant growth occur by way of changes to the infant gut microbiome. We characterize infant gut microbiome organization in three ways, focusing on community diversity, temporal changes in the abundance of microbial taxa, and the sharing of microbiota with maternal reservoirs (i.e. vertical transmission). We further hypothesize that 4) variation in the infant gut microbiome with maternal parity originates in maternal microbial communities, and predict to find that parity effects on the infant gut microbiome are recapitulated in maternal microbial communities.

## RESULTS

### 1. Maternal parity predicts infant postnatal growth

Infant-mother dyads were sampled at three time points across early postnatal life (T1 = 2-5 days old, T2 = 4 months, T3 = 6 months). Despite poorer maternal milk production (Figure S1), infants born to low parity females were larger in body mass at T3 (estimate ± SE: -0.02 ± 0.004, t = -4.51 p < 0.01; Figure S2). Each successive parity was associated with a 0.02 kg decrease in body size at both time points (mean infant body mass at T3 = 1.20 kg). there was no relationship between maternal parity and infant body mass in the first days of life (T1), suggesting that parity-related differences in infant growth emerge primarily across the first 6 months of life and are not influenced by variation in birth weight.

### 2. Parity is associated with infant gut microbiome organization

#### Alpha and beta diversity

Amplicon sequencing of the 16S rRNA gene identified 1,956 unique amplicon sequence variants (ASVs) across all infant gut microbiome samples (mean ± SD = 276 ± 144 ASVs per sample, range = 45-542) compared to 2,019 unique ASVs across maternal gut microbiome samples (mean ± SD = 399 ± 94 ASVs per sample, range = 176-661) and 2,714 unique ASVs across maternal milk samples (mean ± SD = 484 ± 176 ASVs per sample, range = 152-742). Infant gut microbiome alpha-diversity was lower than that of both maternal communities (gut and milk) at T1 (estimate ± SE: -340.43 ± 69.16, t = - 4.92, p < 0.0001) but increased sharply to T2 (estimate +/-SE: 57.00 +/-7.63, t=7.47, *p* < 0.0001; Figure 1A). Compositional differences between the gut microbiome communities of infants and their own mothers (i.e. dyadic beta diversity) were significantly greater at T1 compared to T2 and T3 (estimate ± SE: -0.04 ± 0.007, t = - 6.38, p < 0.0001; Figure S3), suggesting that infants achieve an adult-like gut microbiome by 4 months of age.

**Figure 1.**
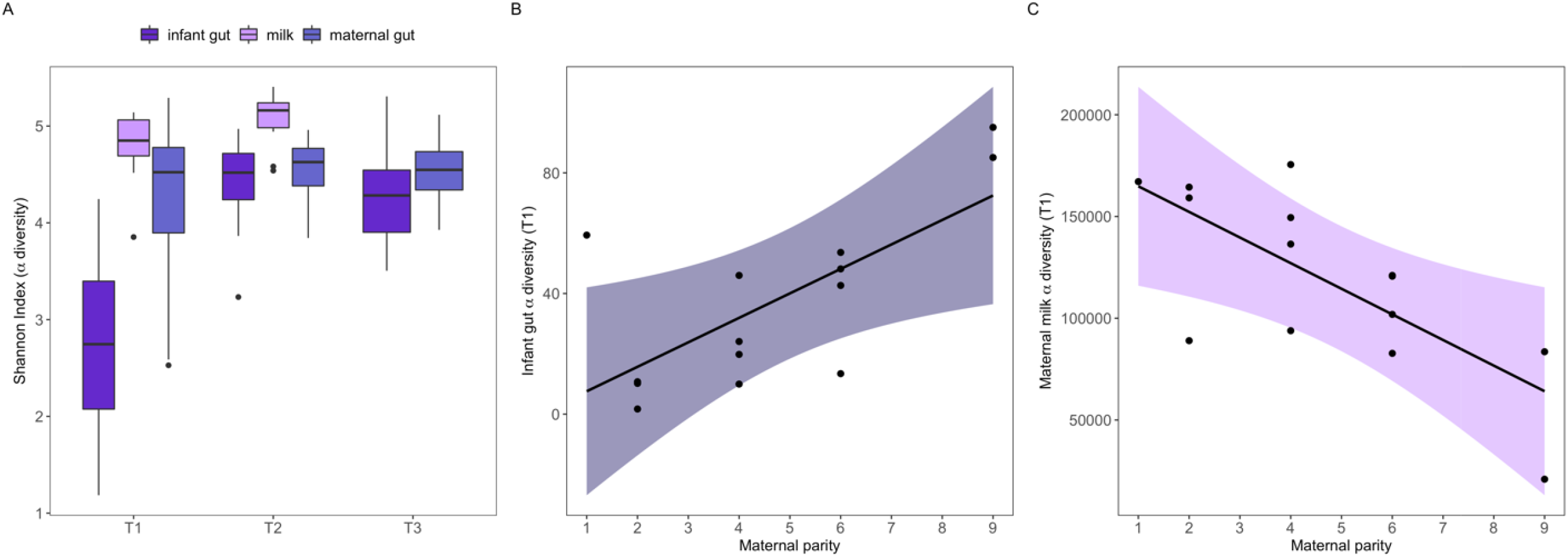
Maternal parity predicts variation in alpha diversity of the infant gut and maternal milk microbiomes. **(A)** The infant gut microbiome exhibits significantly lower Shannon Indices than maternal reservoirs at T1, but converges toward maternal levels by T2. **(B)** Shannon Indices (partial residuals) at T1 are lowest in low parity infants. **(C)** The maternal milk microbiome of low parity females is more diverse (partial residuals of Shannon Indices) than high parity females.

Infants born to low parity females exhibited lower alpha-diversity than high parity infants at T1 (estimate ± SE: 6.13 ± 2.02, t = 3.04, p < 0.01; Figure 1B), but not at T2 or T3. Of the two maternal communities, only the milk microbiome varied with maternal parity, and the pattern was opposite to the infant gut: low parity females exhibited more diverse milk microbiomes than high parity females at T1 (estimate ± SE: -12391 ± 3836, t = -3.23, p < 0.01; Figure 1C), and there was no relationship between maternal microbial alpha diversity and parity at any other time point. There was also no relationship between maternal parity and beta diversity.

#### Differential abundance of taxa

Amplicon sequence variants (ASVs) in the infant gut microbiome were resolved to 78 bacterial families, of which 42 were highly abundant (> 1% of total community composition; Figure S4, Table S1). Of those taxa, 21 families exhibited statistically significant changes in relative abundance from T1 to T2, and 11 families from T2 to T3 (Benjamini-Hochberg adjusted p-values: p_BH_ < 0.05; Table S2). At T1, the infant gut was dominated by a single ASV from the *Bacteroidaceae* family (assigned to *Bacteroides fragilis*; mean relative abundance 28.5%). By T2, infants had almost entirely lost *Bacteroidaceae* from the gut community, which instead became dominated by *Prevotellaceae* (36% relative abundance).

To identify which specific ASVs within these bacterial families contributed the most to infant gut microbiome maturation, we used a Compositional Data Analysis (CoDA) framework to account for the compositional nature inherent to microbiome data [50]. PC1 explained 18.7% of the total variance in composition among infant samples and showed a clear distinction between early life (T1) samples and older samples (T2 and T3) (Figure 2A). Clustering on PC1 was most strongly explained by two ASVs with the highest (absolute value) loading scores: *Bacteroides fragilis* dominance among neonatal samples (T1), and *Prevotella copri* dominance among older (T2 and T3) samples (Figure 2B; Table S2).

**Figure 2.**
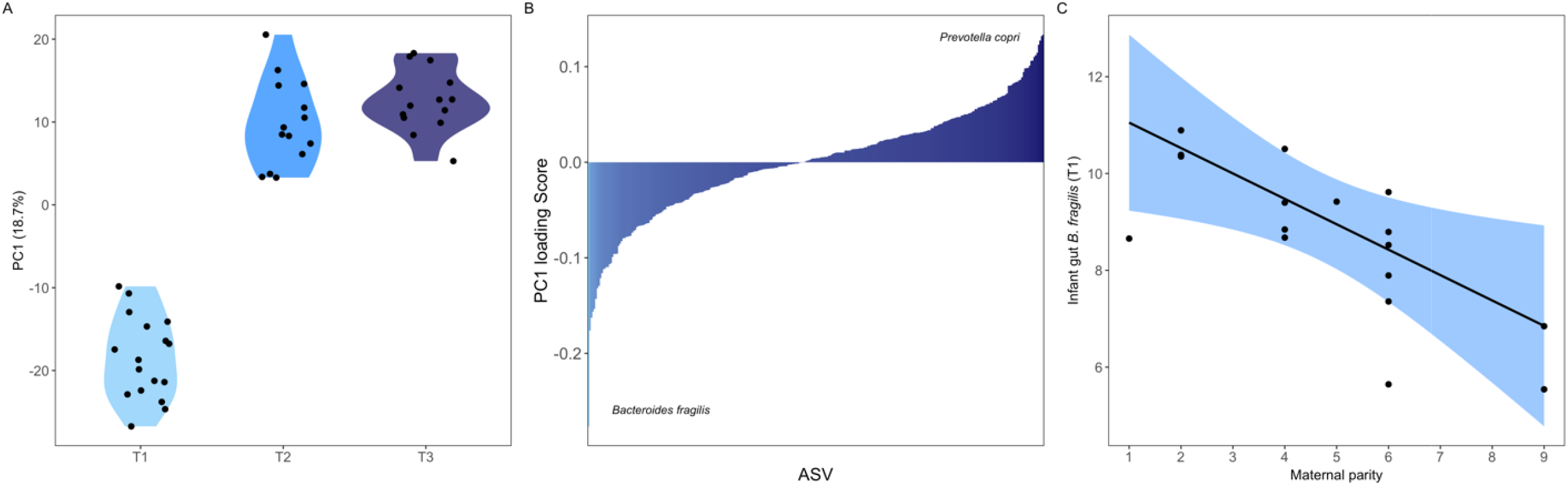
Compositional maturation of the infant gut microbiome is driven by a transition from *Bacteroides fragilis* dominance to *Prevotella* dominance, with infants of low parity females exhibiting more *B. fragilis* during early life. **(A)** Between-sample dissimilarity (Aitchison distance) on the first principal component (PC1) according to infant age. **(B)** Loading scores of all infant gut ASVs on the first principal component, with the most influential ASVs depicted on the left for younger infants (*B. fragilis)* and on the right for older infants (*P. copri)*. **(C)** Partial residual plot demonstrating that infants born to low parity females exhibit significantly higher relative abundances of *B. fragilis* than high parity infants at T1.

Infants born to low parity females exhibited stronger early life dominance of *B. fragilis* compared to high parity infants (estimate ± SE: -0.40 ± 0.14, t = -2.75, p < 0.01; Figure 2C), but there was no relationship between maternal parity and *P. copri*. A greater abundance of *B. fragilis* was significantly associated with lower alpha diversity at T1 (estimate ± SE: -1.15 ± 0.16, t = -7.25, p < 0.001), suggesting that this ASV may be driving microbial community homogeneity during early life.

#### Vertical transmission from maternal reservoirs

At T1, vertical transmission from the milk microbiome to the infant gut was stronger than from the maternal gut microbiome (estimate ± SE: 9.50 ± 4.32, t = 2.20, p < 0.05; Figure 3A). Infants shared 60 ASVs on average with their mother’s milk microbiome (51% transmission rate) versus 41 ASVs with their mother’s gut microbiome (42% transmission rate). ASV sharing between the infant gut and maternal milk microbiome was greater in low parity dyads than high parity dyads (estimate ± SE: -7.51 ± 2.03, t = - 3.69, p < 0.01; Figure 3B). There was no effect of maternal parity on infant ASVs shared with the maternal gut microbiome. By T2, the average transmission rates from milk and the maternal gut to infants were not significantly different, and there was no longer an effect of maternal parity on transmission rates from milk.

**Figure 3.**
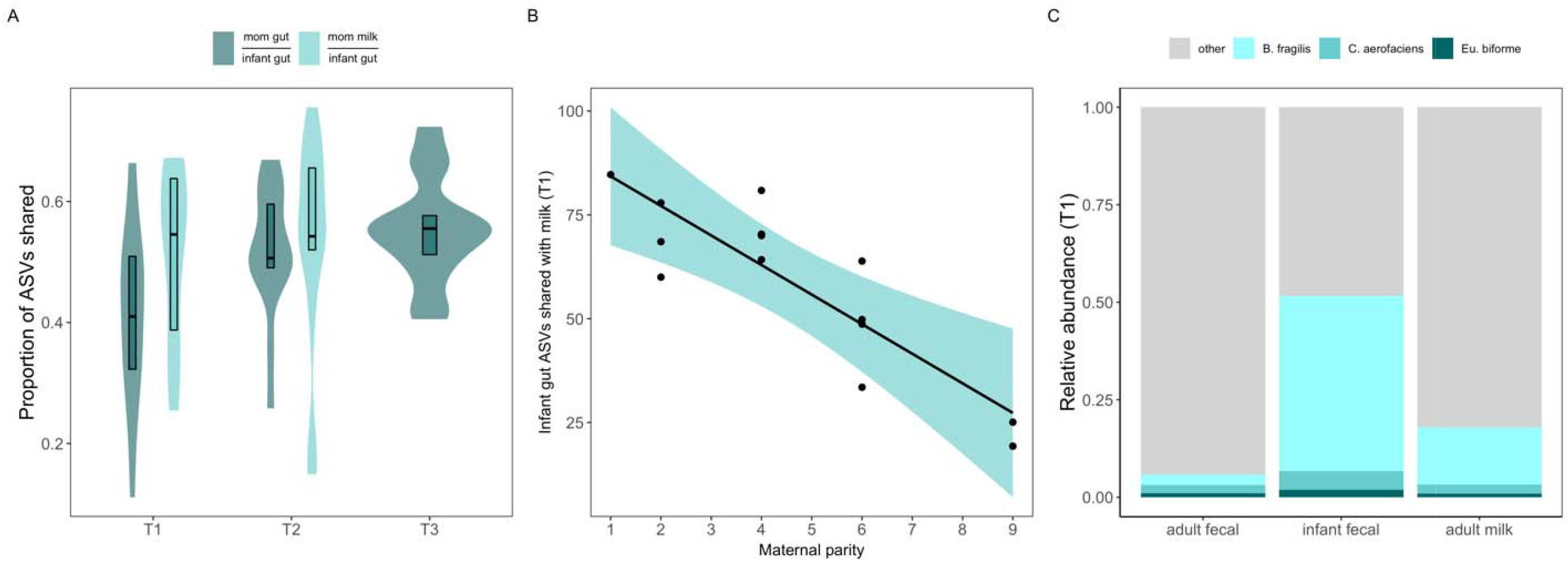
Vertical transmission via the milk microbiome varies with maternal parity. (A) Across all infants, the infant gut microbiome shares ∼10% more ASVs with milk than with the maternal gut at T1. Transmission rates do not differ at T2. (B) Infants born to low parity females exhibit stronger ASV sharing with the milk microbiome at T1 compared to infants of high parity females. (C) Average relative abundances of the 3 high-frequency ASVs across microbial communities suggests that milk is the primary reservoir for *B. fragilis* to the infant gut.

To pinpoint which shared ASVs may be important for infant development, we identified ASVs that were both highly prevalent (i.e. present in all infant gut microbiome samples in our dataset) and shared at high frequency with the maternal milk and gut communities (i.e. present in matched mother-infant samples across ⍰90% of dyads). Three ASVs were prevalent and shared at high frequencies: *Bacteroides fragilis, Collinsella aerofaciens*, and *Eubacteria biforme. B. fragilis* and *Eu. biforme* were shared exclusively between maternal milk and infant gut microbiomes, while *C. aerofaciens* was shared universally across all three communities. The relative abundances of *C. aerofaciens* and *Eu. biforme* were similar across maternal and infant communities (Figure 3C). However, the infant gut housed by far the greatest proportion of *B. fragilis* (Figure 3C). Mothers housed ∼15x more *B. fragilis* in their milk compared to their gut (Figure 3C), evidence that infant gut *B. fragilis* may originate in milk.

There was no significant relationship between maternal parity and the relative abundance of *C. aerofaciens* or *Eu. biforme* in the infant gut microbiome, nor was there a relationship between maternal parity and these two microbial taxa in the maternal gut microbiome. However, low parity females housed significantly more *Eu. biforme* (estimate ± SE: -0.09 ± 0.04, t = -2.47, p < 0.05) and *B. fragilis* (estimate ± SE: -0.09 ± 0.04, t = -2.01, p < 0.05) in their milk compared to high parity females.

### 3. The early life infant gut microbiome mediates parity effects on postnatal growth

We constructed three path analyses corresponding to the three parity-dependent measures of infant gut microbiome composition at T1 (alpha diversity, *Bacteroides fragilis* abundance, and ASVs shared with the maternal milk microbiome) to test whether parity effects on infant body mass at T3 were mediated by the early infant gut microbiome. Path models revealed that the effects of maternal parity on infant body mass were mediated by differences in infant gut microbiome organization, as the direct effect of parity of infant mass became non-significant. There was no mediation effect of ASVs shared with the milk microbiome.

In the diversity path model, the effect of maternal parity on neonatal infant gut alpha diversity was the strongest path present (***β*** = 0.709, p < 0.01), and parity indirectly influenced infant mass via reduced infant gut alpha diversity (***β*** = -0.269, p < 0.05) (Figure 4A-B). Infants with the greatest body mass also possessed the least diverse gut microbiomes. A reduction in each unit of early life Shannon Diversity was associated with a gain of approximately 0.3 kg in later-life body mass. Similarly, in the *B. fragilis* model, maternal parity influenced infant mass via relative abundance of *B. fragilis* in the infant gut (***β*** = 0.359) (Figure 4C-D). With each unit increase in the relative abundance of early life *B. fragilis*, infants gained approximately 0.4 kg in body mass.

**Figure 4.**
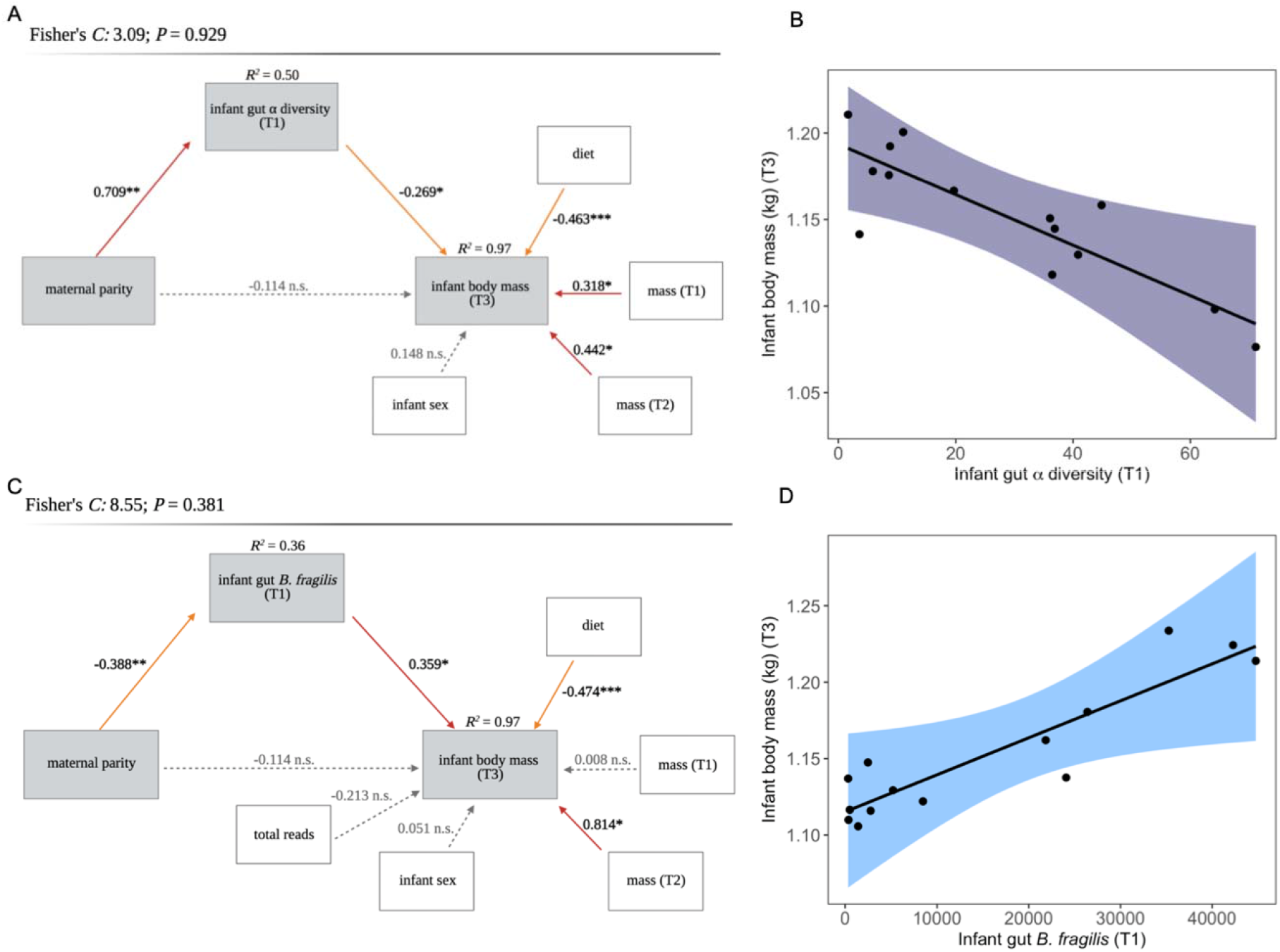
A specialized early life infant gut microbiome abundant in *B. fragilis* mediates the effects of maternal parity on infant growth. Path models showing the indirect effects of maternal parity on infant body mass at 6 months old (T1) via early life infant gut **(A)** alpha diversity and **(B)** *B. fragilis* relative abundance. Global goodness-of-fit was evaluated using Shipley’s test of d-separation (*P* ≥ 0.05 indicates a good fit) and is reported at the top of each path model. Variables of interest are housed in grey boxes, while control/confounding variables are in white boxes. Solid red arrows represent significant positive paths, solid orange arrows represent significant negative paths (*P* ≤ 0.05), and grey dashed arrows indicate non-significant paths. Asterisks denote significant paths (*P* ≤ 0.05 *, *P* ≤ 0.01 **, *P* ≤ 0.001 ***). *R*^*2*^s of each of the component models are located on top of the corresponding endogenous (dependent) variables. Partial residual plots of component models additionally highlight the mediation effects of infant gut alpha diversity **(C)** and *B. fragilis* relative abundance **(D)** on infant body mass, independent of maternal parity.

## DISCUSSION

The data presented here demonstrate that the early life infant gut microbiome mediates the relationship between maternal parity and infant growth. Despite poorer milk production, infants born to low parity females attained greater body mass by 6 months of age and exhibited divergent patterns of microbiome organization. Lower alpha diversity and more *B. fragilis*, but not stronger vertical transmission, mediated parity effects on infant growth. Further, among maternal communities, parity effects were only apparent in the milk microbiome, indicating that it may be the source of parity-dependent variation in the infant gut. Together, our findings suggest that the first days of life are a critical period of infant gut microbiome organization through which maternal condition shapes the pace of postnatal growth.

Consistent with previous studies in humans and rhesus macaques but opposite a recent study in chimpanzees [51,52], alpha diversity of the vervet monkey infant gut microbiome was lowest during early life, converging rapidly toward maternal gut microbiome levels by 4 months old. Although a less diverse microbiome during adulthood can indicate microbiome dysbiosis [53,54], reduced diversity during development may instead reflect specialization of the gut microbiome for specific functions. Indeed, the infant gut microbiome of breastfed human infants is less diverse than formula-fed infants and houses more microbial taxa involved in milk glycan metabolism [55–57]. Increased metabolism of milk glycans, which hosts cannot digest on their own, prevents pathogen invasion by modifying the resource landscape in the gut [58] while also providing energetic resources to fuel host growth and development [25]. The reduced alpha diversity characteristic of early life, particularly in infants born to low parity females, may reflect a taxonomically and functionally homogeneous microbial community, perhaps to better utilize smaller volumes of milk [57]. In support of this hypothesis, low alpha diversity at T1 predicted greater body mass at T3 in vervets, independent of maternal parity. These results are in line with studies on birds showing an association between reduced early life alpha diversity on future weight gain [23,24]. The first days of life thus appear to represent a critical period of microbiome organization in which community diversity can impact later-life host growth.

While there was no relationship between maternal parity and alpha diversity of the maternal gut microbiome, low parity females exhibited a more diverse milk microbiome than high parity females. Interestingly, this relationship was opposite that of the infant gut (infants: low parity/high alpha diversity, mothers: low parity/low alpha diversity). This contradiction suggests that the reduced diversity of the low parity infant gut microbiome is not simply a result of transmission from a less diverse maternal milk microbiome. Instead, this pattern provides evidence for stronger selective seeding from a diverse milk microbiome in low parity infants [59].

At the taxonomic level, the infant gut microbiome changed more dramatically from T1 to T2 than from T2 to T3. At T1, the infant gut was dominated by *Bacteroidaceae*, but shifted to *Prevotellaceae* dominance by T2. At the ASV level, changes in infant gut microbiome composition with age were most strongly explained by two ASVs: *Bacteroides fragilis* and *Prevotella copri*. A trade-off between *Bacteroides* and *Prevotella* has been well-described in microbiome research for over a decade as a reflection of dietary differences between human populations consuming a diet rich in fats and proteins (more *Bacteroides*) and a diet rich in fiber (more *Prevotella*) [60,61]. In young vervet monkeys, a trade-off between these taxa may serve a developmental role as infants transition from the consumption of milk to solid foods. Unlike *B. fragilis*’s role in milk glycan consumption [26,62], *P. copri* cannot digest glycans [60], which comprise a significant proportion of human and nonhuman primate milk [63,64]. Indeed, nursing piglets exhibit greater *Bacteroidaceae*, while weaned piglets exhibit greater *Prevotellaceae* [65].

Across maternal parities, infants appear to selectively seed *B. fragilis* in the first days of life (T1), with greater abundances of *B. fragilis* predicting greater body mass at T3, independent of parity. However, infants of low parity females exhibited greater *B. fragilis* abundance than their counterparts, potentially indicating stronger selection for this particular microbe. Low parity females can produce as little as 5x less milk than high parity females in this population (Figure S1). Stronger selection for *B. fragilis* among low parity infants may thus serve to enhance host digestion of glycans from lower volumes of milk, resulting in a microbial community that is more efficient at assimilation milk components to fuel growth. In support of this explanation, experimental studies on rodent models of the human microbiome have found that greater abundance of *B. fragilis* promotes infant growth in the presence of milk glycans [25]. Moreover, *B. fragilis* dominance appears to be the driver of infant gut microbiome specialization at the community level: a greater abundance of early life *B. fragilis* predicted overall lower gut microbiome alpha diversity at both T1 and T2. Island biogeography theory predicts that early colonizing bacteria can impact the trajectory of gut microbiome development by exerting influence over immigration order through historic ecological processes [59]. *B. fragilis* modulates the biochemical makeup of the gut environment [62,66], which in turn can alter subsequent colonization by other taxa. Though largely absent from the infant gut microbiome at T2, *B. fragilis* dominance at T1 may explain the persistence of reduced community diversity via regulation of immigration order [26,62].

The sharing of amplicon sequence variants has been previously used as evidence of vertical transmission from maternal reservoirs to the offspring [37,45,67,68]. At T1, vervet infants shared significantly more (∼10%) ASVs with their own mother’s milk microbiome than with her gut microbiome, resulting in a transmission rate from milk of ∼51% compared to ∼42% from the maternal gut. These rates of transmission are overall higher than previously reported transmission rates from milk [44] and the maternal gut [28] in humans. Higher rates may reflect stronger vertical transmission via milk in nonhuman primates compared to humans, and suggests that transmission and the incorporation of milk microbiota into the infant gut is under strong evolutionary selection [59]. Notably, our findings contrast with previous studies on humans that suggest that the maternal gut microbiome is the largest source of colonizing taxa to the infant gut [28,35,69], thus calling for the inclusion of milk as a maternal reservoir in comparative studies on maternal transmission.

While there was no relationship between maternal parity and ASV sharing with the maternal gut microbiome at any age, infants of low parity females shared significantly more ASVs with their mother’s milk microbiome than high parity infants at T1. There are a few potential explanations for this finding: first, the dispersal of milk microbiota may be enhanced in low parity females. The vervet monkey milk microbiome is a highly individualized community that houses many “private”, or individualized ASVs [41]. Low parity females may possess more individualized variants that are better adapted for dispersal to and survival within the infant gut [70], an explanation also supported by our finding of greater taxonomic diversity in the milk microbiome of low parity females.

Second, low parity females may exhibit variation in non-microbial components of milk (e.g. oligosaccharides, immunoglobulin A), which can modulate the infant gut environment in a manner that favors colonization by maternal milk ASVs over other taxa [71,72]. Finally, low parity infants may selectively seed maternal milk microbiota more strongly than high parity infants via maternally-directed, or infant-specific mechanisms that promote colonization by maternal milk ASVs (e.g. phage activity, mucosal immunity, pH) [59].

High frequency sharing of specific microbial variants provides evidence that the selective seeding of these taxa may be evolutionarily conserved [29]. At T1, there was a 15-fold higher abundance of *B. fragilis* in the milk microbiome compared to the maternal gut microbiome, suggesting that milk may be the primary reservoir of *B. fragilis* to the infant gut [47]. *B. fragilis* relative abundance was significantly greater in the milk microbiome of low parity females compared to high parity females, indicating that the milk microbiome may be the source of parity-dependent differences in *B. fragilis* colonization in the infant gut. Moreover, given that neither the maternal vaginal nor gut microbiome explain variation in infant *Bacteroides* colonization in prior studies [33,47], our data suggest that milk may also be the primary source of *B. fragilis* more broadly. Based on these interpretations, we encourage future research on the infant gut microbiome to consider the importance of the milk microbiome and milk *B. fragilis* for early infant development.

This study was limited in a couple of important ways. First, although the vaginal microbiome (an additional major maternal reservoir of taxa to the infant gut) may contribute to or modify the above-described relationships, logistic constraints prevented vaginal microbiome sampling in this study. Despite this limitation, prior studies in humans indicate that the vaginal microbiome is unlikely to be the primary contributor of *B. fragilis* to the infant gut [44]. Given the importance of *B. fragilis* in mediating parity effects on infant growth in this population, our interpretation of milk as the potential source of *B. fragilis* is likely independent from vaginal microbiome effects. Second, the correlation between maternal parity and maternal age raises the question of whether the relationships described here are simply age effects. As our hypotheses were centered on the effects of successive reproductions on maternal communities, in particular the milk microbiome, we did not include maternal age in our analyses to avoid multicollinearity. Effects of parity, rather than age, on microbiome composition may be unsurprising for two reasons: successive reproductions independent of maternal age shape the morphology of the mammary gland [46] and variation in milk production across mammalian species (e.g. [13,14]), and age does not typically exert strong effects on microbiome composition in mammals.

Viability selection exerts substantial pressure on infant development in populations with high rates of early mortality [73,74]. In our study population, infants face a growth-associated mortality rate of more than 30% [48], whereby individuals who die early in life are smaller in body size than their counterparts [49]. Given these pressures, our data suggest that low parity females may compensate for the costs of poor milk production by modifying, potentially through alterations to the milk microbiome, transmission and seeding to favor the establishment of a more specialized, milk-oriented infant gut microbiome. As greater microbial specialization for the consumption of milk components is broadly adaptive for mammalian infants [56], our findings suggest that reduced alpha diversity, and more *B. fragilis*, during early life may be adaptive in populations at high risk of growth-associated infant mortality.

## CONCLUSION

As in human infants, *Bacteroides fragilis* plays a crucial role in establishing a specialized, milk-oriented microbiome in infant vervet monkeys. Enhancing *B. fragilis* colonization in low parity infants, via stronger selective seeding or vertical transmission, may serve as a compensatory mechanism by which infants born to low parity females can enhance intestinal assimilation of milk components from lower volumes of milk. Our study adds to a growing body of literature on maternal effects on infant gut microbiome composition and development broadly, and is unique in its incorporation of the milk microbiome. We encourage future studies on the developmental microbiome in mammals to consider the potential role of the milk microbiome in driving successional patterns in the infant gut as well as influencing postnatal growth.

## METHODS

### Study population

Subjects for this study were 18 vervet monkey mother-infant dyads housed at the Vervet Research Colony (VRC) at the Wake Forest School of Medicine, Winston-Salem, North Carolina. Vervets are highly social primates that breed once annually [75].

Mother-offspring dyads were housed at the VRC across 8 indoor-outdoor social groups organized by female kin. Thirteen of the dyads were provisioned daily with commercial monkey chow (Purina Monkey Chow, LabDiet 5038), while four dyads were fed a western-style lab diet (LabDiet 5L3K) as part of a different study. In addition, some dyads were fed ad libitum, while feeding was time-restricted for others. However, it is important to note that infants in this population do not appear to incorporate solid foods into their diets until about 4-6 months [76], thus the effect of diet on infant microbiome composition in this study is expected to be minimal. Animals were supplemented with fresh fruits and vegetables daily. Infants included in this study were delivered via unassisted vaginal births between June 18, 2017 and August 27, 2017 after 5.5 month gestations.

### Sample collection

We collected physiological data from dyads at three postnatal timepoints: T1 (2-5 days postpartum), T2 (4 months postpartum), and T3 (6 months postpartum). Two infants died within the first month of life as a probable consequence of poor maternal milk production: in one case, the mother was not producing any milk, and in the second case, the mother produced milk in only one mammary gland. Thus both infants contributed fecal samples and somatometric data at T1 only, and the T1 milk sample from the female with unilateral milk production was excluded from this study. Milk collection followed previously published protocols for the collection of milk from cercopithecine monkeys [17,41]. Both mammary glands were fully evacuated via manual expression into a single sample tube. Samples were placed immediately on ice, briefly vortexed, aliquoted into cryovials, and frozen at −80°C until shipment to the University of Washington. We collected maternal and infant fecal samples at all three sampling timepoints by briefly inserting standard (mothers) and pediatric (infants) flocked nylon swabs (FLOQSwabs, COPAN Diagnostics, Murrieta, CA) into the anal canal. Rectal swabs are reliable proxies for fecal samples when quantifying an animal’s gut microbiome, including among infants [77,78]. Swabs were gently spun in the canal 2-3 times before removal and were snapped off into empty polypropylene tubes. Samples were immediately frozen at −80°C before being shipped to the University of Washington.

### DNA extraction, amplification, and library preparation

We extracted microbial DNA from milk using the PowerFood Microbial kit (Qiagen) following the manufacturer’s kit protocols, with the addition of two front-end processing steps shown to increase overall yield from milk [41]. For fecal swabs, microbial DNA was extracted using the Qiagen PowerLyzer PowerSoil kit. Samples were thawed to room temperature, swab tips were snapped into glass bead tubes (0.1 mm), and mechanically lysed (30 Hz for 2 min). Extractions then followed the manufacturer’s protocol.

We amplified the hypervariable V4 region of the 16S rRNA gene using PCR primer set 515F and 806R from The Human Microbiome Project [79,80], following previously published protocols [41]. A fragment analyzer was used to confirm amplification of the V4 region prior to quantification via a qubit fluorometer.

### Sequencing and bioinformatics

Amplicon libraries were balanced and pooled. Library complexity was increased by spiking libraries with PhiX prior to sequencing all libraries together on a single Illumina MiSeq flow cell using 301□bp paired-end sequences. This resulted in 1,456,972 reads for milk samples, 2,491,940 reads for adult fecal samples, and 2,147,458 reads for infant fecal samples. Sequences were analyzed using the Quantitative Insights Into Microbial Ecology 2 (QIIME2) platform [81,82] and denoised with Divisive Amplicon Denoising Algorithm 2 (DADA2:[83]) as a QIIME2 plug-in. In contrast to clustering sequencing reads based on a fixed dissimilarity threshold (e.g., the assignment of Operational Taxonomic Units [OTUs]), which can conflate sequencing errors with biological variation, DADA2 infers sequences exactly (resulting in amplicon sequence variants, hereafter referred to as ASVs), providing higher taxonomic resolution than OTUs [83,84]. This approach is crucial for assessing changes in microbial composition [85] as well as community diversity [86], particularly in microbial communities presumed to be low in diversity based on prior OTU clustering methods (e.g., the human vaginal microbiome [84]), as well as determining vertical transmission between maternal and infant communities. Forward and reverse reads were trimmed to 240 bases long to remove the low-quality portion of the sequences. Next, the forward and reverse reads were merged and chimeric sequences removed. After filtering, trimming, merging, and chimera removal, we retained 811,225 16S rRNA gene sequences from 29 milk samples (28,972□±□17,165 reads per sample), 1,547,442 sequences from 45 adult female fecal samples (48,357□±□15,100 reads per sample), and 2,147,458 sequences from 45 infant fecal samples (47,721 ± 12,580 reads per sample). Only fecal samples with >15,000 reads were retained for analyses, resulting in N=44 infant fecal samples. No other samples were removed at this stage as sequencing depths across samples were highly balanced (Figure S5). ASVs were aligned using MAFFT [87] and a phylogenetic tree was constructed using fasttree2 [88]. Taxonomic assignment of ASVs was performed using the q2-feature-classifier in QIIME2 against the 13_8 version of the GreenGenes database [89] based on 100% similarity to the reference sequence. One milk sample was excluded from the analysis as all of its 200,000+ reads were assigned to a single ASV.

### Statistical analyses

All analyses for this study were performed in QIIME2 [82] and R version 4.0.2 [90]. Taxonomy and count tables were imported from QIIME2 into R using the qiime2R package [91]. As rarefaction of microbiome count data results in a loss of precision and variation [92], we instead used a compositional data, or “CoDa”, approach when necessary [50] and controlled for sequencing depth by including the log-transformed count of total reads in a sample as an offset in all statistical models. In addition, all statistical models constructed in this study included diet as a fixed factor. Where microbial taxa were converted to relative abundances, the taxonomy table was filtered to retain only those with > 1% relative abundance. Residuals of all linear models described below were found to be normally distributed following visual evaluation by Q-Q plots and Shapiro-Wilk tests. Because of the potential confounding effects of dietary composition and schedule, diet was controlled for as a fixed factor in all analyses. Figures were created in R using ggplot2 [93] and visreg for partial residual plots [94].

#### Milk yield

Linear models were used to test the effects of maternal parity on milk production (mL). Residuals for all models were normally distributed upon visual inspection and evaluation with a Shapiro-Wilk test, and therefore not transformed prior to analysis. We constructed models with milk yield (mL) as the dependent variable, with maternal parity as a fixed factor while controlling for offspring sex [17,95] and dietary composition.

#### Infant body mass

Linear models were used to test whether maternal parity predicted infant body mass at T3 (dependent variable), controlling for infant sex, approximate birth weight (body mass at T1), and dietary category. T3 body mass data were log-transformed prior to modelling to ensure normality of the model residuals.

#### Alpha & beta diversity

Microbiome samples were assessed at the community level using both alpha (within sample) and beta (between samples) diversity measures. Alpha diversity was calculated using the phyloseq package [96] and quantified using the Shannon Index of alpha-diversity [97,98], which takes into account both the richness and evenness of taxa. Shannon Indices were transformed by Tukey transformation to achieve normality of residuals, and linear mixed models were constructed to test associations between infant age (categorical fixed effect: T1, T2, T3) and gut microbiome diversity (Shannon Index, dependent variable), controlling for diet (fixed effect) and infant ID (random effect). Weighted UniFrac distance matrices were constructed in QIIME2 and were used to assess dissimilarity between the gut microbiome of infants and their own mothers (dyadic weighted UniFrac distance) as a measure of infant gut microbiome maturation. A linear mixed model was used to test whether infant age (fixed effect) predicted dyadic weighted UniFrac distance (dependent variable), controlling for diet (fixed effect) and infant ID (random effect).

#### Differential abundance testing

Differential abundance testing of taxa within the infant gut microbiome was performed in R using the NBZIMM package [99]. Negative binomial models (glmer.nb() function) were used test the effects of infant age (fixed effect) on the abundance of taxonomic families (dependent variable: raw counts), controlling for diet (fixed effect), sequencing depth fit as an offset (log transformation of total number of reads in the sample), and infant ID (random effect). Models that did not converge due to a high proportion of zeros were rerun using the same package as zero-inflated negative binomial models (lme.zig() function). P-values were adjusted using the Benjamini-Hochberg FDR multiple-test correction [100] to account for multiple hypothesis testing and microbial taxa with adjusted p-values < 0.05 were considered statistically significant.

#### Principal Components Analysis & CoDa framework

A Compositional Data (CoDa) framework was used to investigate how specific ASVs contributed to the compositional patterns of infant gut microbiome maturation [50]. Raw count data of ASV reads were normalized using a centered log-ratio (clr) transformation and a pseudocount of 0.65 in place of zeroes [101]. We then ordinated the samples based on their compositional dissimilarity using a Principal Components Analysis (PCA) [102] using pairwise Euclidean distances between samples (i.e. Aitchison distance). Unlike Principal Coordinates Analysis (PCoA), which is more commonly used in microbiome data analysis, PCA is not driven by presence/absence data, is more reproducible, and is more robust against sparsity [50]. The PCA was used to visualize sample clustering and generate loading scores for each ASV on the first PCA axis (which captured 18.7% of variability and was highly predicted by infant age). The loading scores were then sorted by absolute value to determine the microbial taxa with the highest influence (e.g. absolute value loading score) on the observed clustering of the first PCA axis (i.e. the most predictive of infant age) [103]. We then calculated a log-ratio of the relative abundance of the two most influential taxa (*Bacteroides fragilis* and *Prevotella copri*) and used Pearson’s correlation tests to determine the correlation coefficient between the log-ratios and samples’ coordinates on the first PCA axis.

Subsequently, we used a linear mixed model to test whether the log-ratio of these taxa (fixed effect) predicted dyadic weighted UniFrac distances between infant and maternal gut microbiomes (dependent variable), controlling for infant age (fixed factor), diet (fixed factor), and dyad ID (random).

#### Vertical transmission

The proportion of ASVs shared between maternal and offspring communities is a commonly-used index of vertical transmission using 16S rRNA gene data [29,37,67,68]. We calculated the proportion of shared ASVs as being the proportion of all ASVs present within the infant gut microbiome (denominator) that were also found in maternal milk and gut microbiomes (numerator). To test whether these proportions differed significantly between infant gut/maternal gut versus infant gut/maternal milk comparisons, we fit linear mixed models with proportion of ASVs shared at T1 and T2 as the dependent variable and comparison (milk vs. infant gut or maternal gut vs. infant gut) as a predictor variable, controlling for individual ID. To test whether maternal parity affected vertical transmission, generalized linear models were used to test whether maternal parity predicted the raw count of shared ASVs (dependent variable), controlling for diet and the total number of ASVs present within the infant gut.

#### Mediation analysis

To test for a mediation effect of the microbiome on infant growth, we performed path analyses by constructing three separate path models (one for each parity-dependent measure of infant gut microbiome composition: alpha diversity, *B. fragilis* abundance, and vertical transmission via the milk microbiome) using the piecewiseSEM() in R (v. 2.0.1) [104]. Unlike simple structural equation modeling, piecewiseSEM is robust to small sample sizes and can integrate mixed effects and generalized linear models (e.g. negative binomial models) [104], making it more suitable for use with microbiome count data. Each path model was comprised of two recursive component models: 1) a model testing direct/indirect effects of both maternal parity and the relevant measure of infant gut microbiome composition at T1 on infant body mass at T3, controlling for confounding variables, and 2) a model testing the effect of maternal parity on the infant gut microbiome, controlling for confounding variables. Component models were integrated into a single causal network using the psem() function, which produced weighted coefficients and corresponding P-values to permit the assessment of the significance and relative strengths of each path within the overall network. The global goodness-of-fit was assessed for each model in piecewiseSEM() using Shipley’s test of d-separation [105]. All models had Fisher’s C statistics with P > 0.05, indicating that the models were suitable fits and no major paths were missing.

## DECLARATIONS

### Consent for publication

Not applicable.

### Availability of data and materials

16S rRNA gene sequences will be available at the time of publication under BioProject ID PRJNA728247 at the NCBI Sequence Read Archive (SRA)

(www.ncbi.nlm.nih.gov/sra/PRJNA728247). Data and code for this project are available on figshare (https://figshare.com/s/79f3cf6cf410f5a87d31).

### Authors’ contributions

Concept, design, and interpretation of data: L.P., N.S.M., A.L.; data acquisition and methodology: L.P., M.J.J., S.S., N.S.M., A.L.; writing of original manuscript draft: L.P., A.L.; review and editing of manuscript: A.B., M.J.J., S.S., N.S.M.; funding: L.P., M.J.J., N.S.M., A.L.; supervision: A.L. All authors have approved this version of the manuscript.

## Acknowledgements

We would like to thank the veterinary and technical staff of the Vervet Research Colony, especially Edison Floyd and Chrissy Long, for their role in sample collections, and Katie Hinde for providing expertise and protocol guidance for milk sample collection. We would also like to thank Andreas Koenig and Carola Borries for comments on earlier versions of this manuscript.

