## Supplementary figures legend for "Early life gut microbiome dynamics mediate maternal effects on infant growth in vervet monkeys"

**Supplemental figure 1**. Low parity females produce significantly lower milk volumes than high parity females at T1 **(A)** and T2 **(B).**

**Supplemental figure 2.** Infants born to low parity females are significantly larger than infants born to high parity females T3. Partial residual plot shows the relationship between maternal parity and infant body mass (kg) (T1)**,** controlling for diet, infant sex, and neonatal body mass.

**Supplemental figure 3.** Compositional dissimilarity between the infant gut microbiome and the gut microbiome of their own mothers decreases sharply by 4 months of age (T2). Violin plot of dyadic weighted Unifrac distances across sampling time point, controlling for diet.

**Supplemental figure 4.** Ecologically-oriented heatmap of the 42 most abundant bacterial families in the infant gut microbiome.

**Supplemental figure 5. (A)** Sequencing depth and **(B)** number of unique ASVs by sample type.
