## Supplementary tables legend for "Early life gut microbiome dynamics mediate maternal effects on infant growth in vervet monkeys"

**Supplemental table 1.** Relative abundance of the 42 most abundant bacterial families in the infant gut microbiome through six months of age.

**Supplemental table 2.** Differentially abundant bacterial families from T1 to T2 and T2 to T3. Results, including FDR-adjusted P values, from negative binomial, zero-inflated negative binomial, and linear mixed models.

**Supplemental table 3.** Loading scores of infant gut microbiome ASVs on the first principal component. Negative loading scores correspond to younger infant samples, positive loading scores correspond to older infant samples.
