## Supplementary figures and images for "Early life gut microbiome dynamics mediate maternal effects on infant growth in vervet monkeys"

### Supplementary figure 1

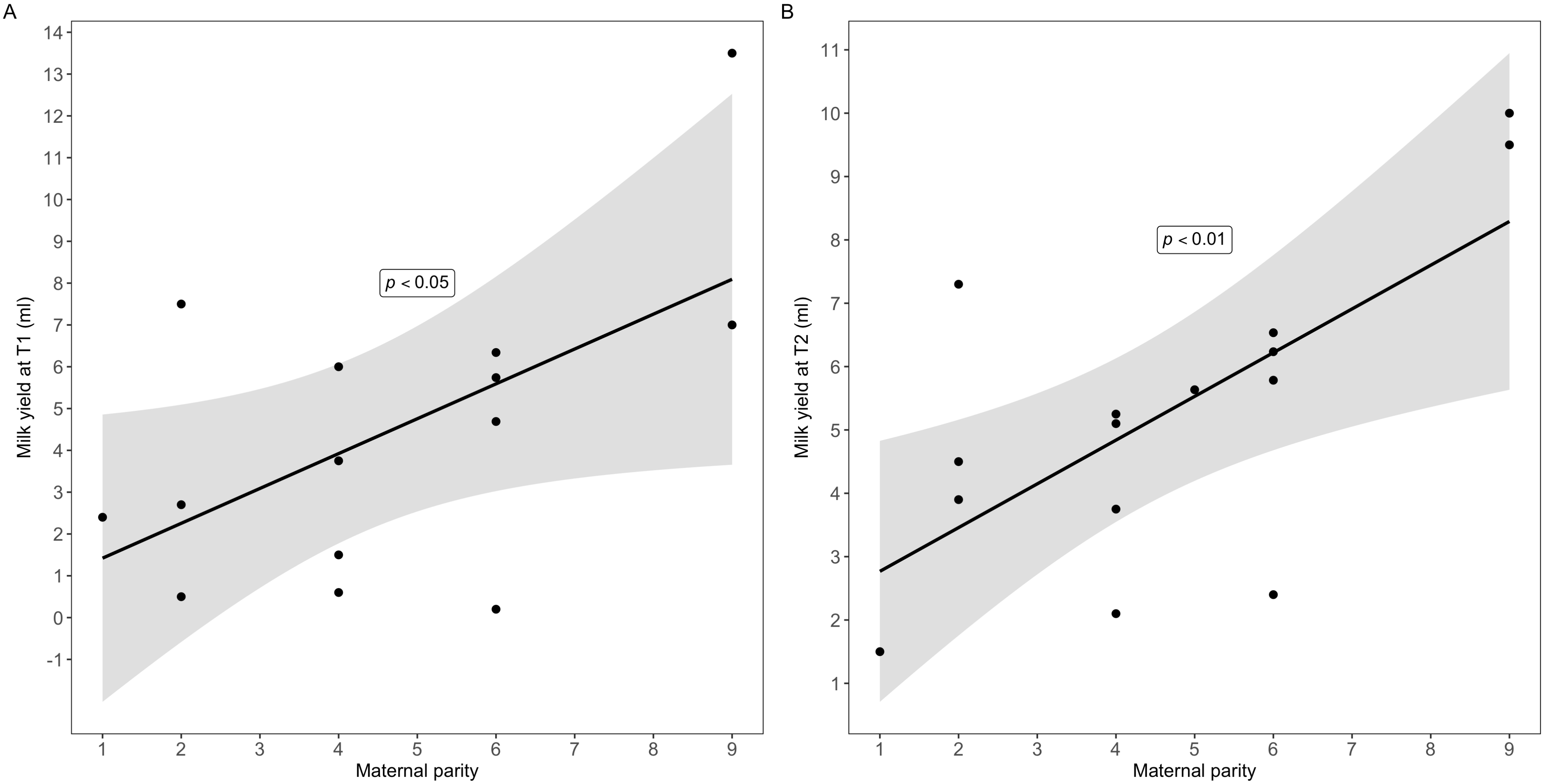

### Supplementary figure 2

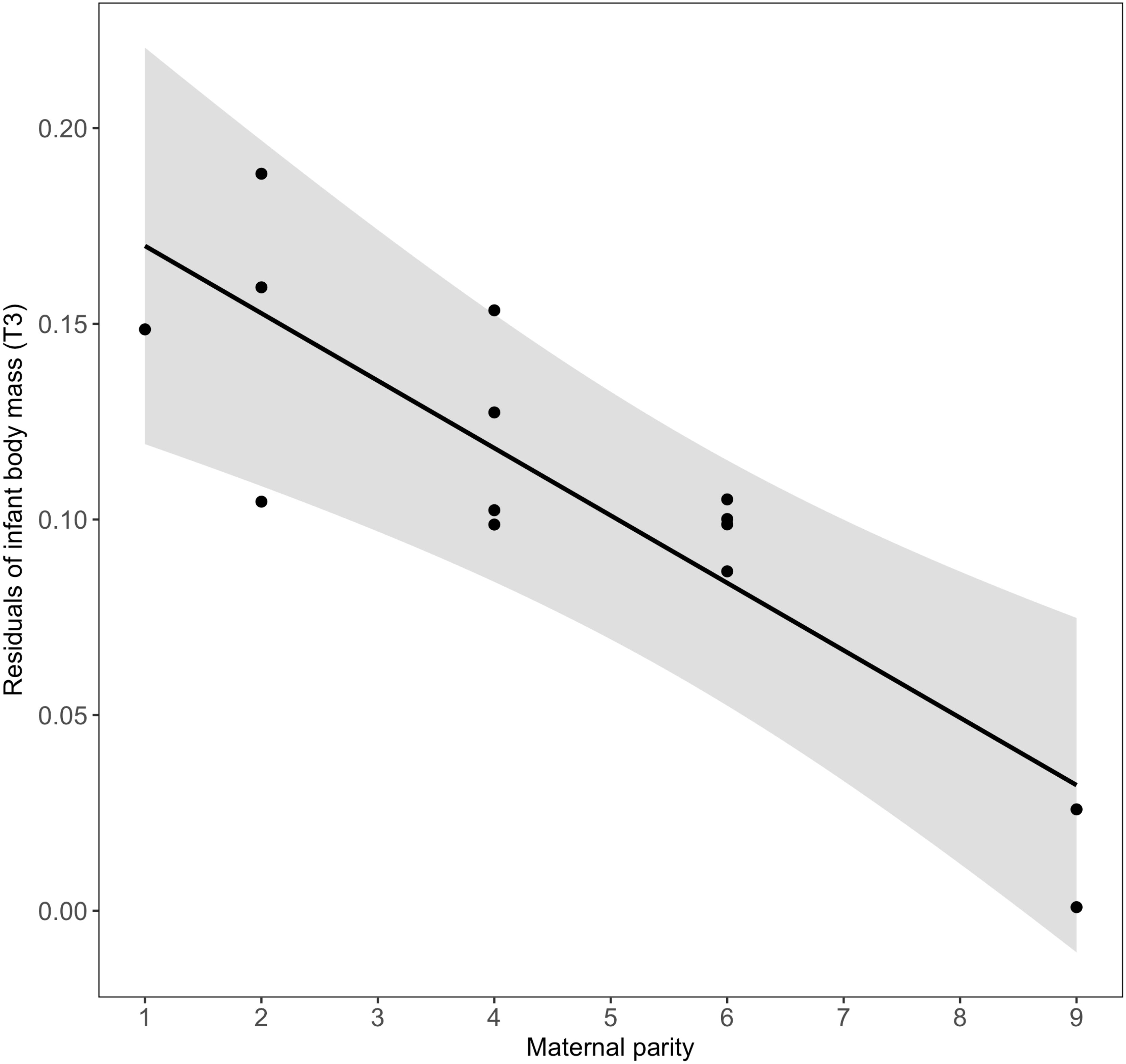

### Supplementary figure 3

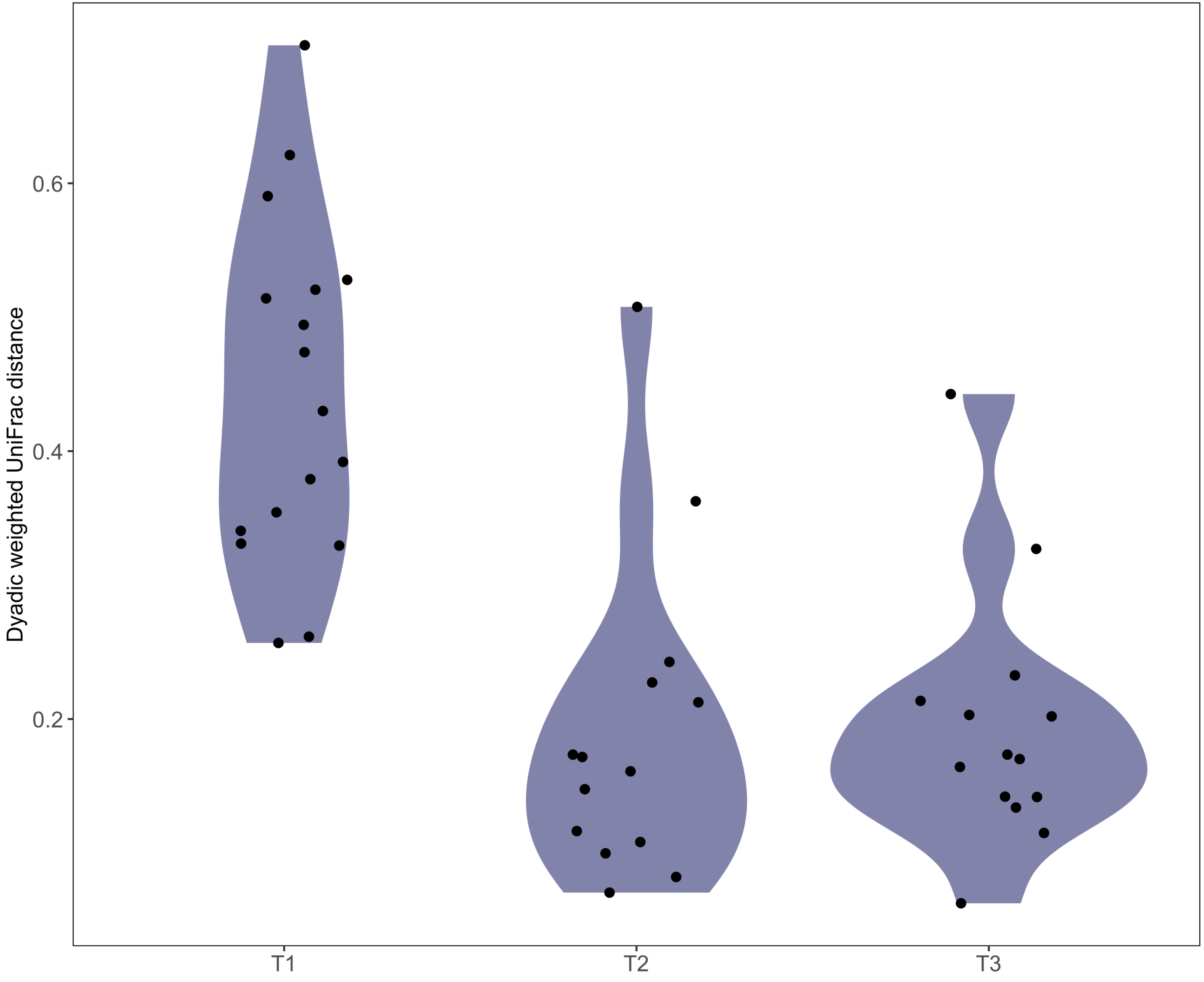

### Supplementary figure 4

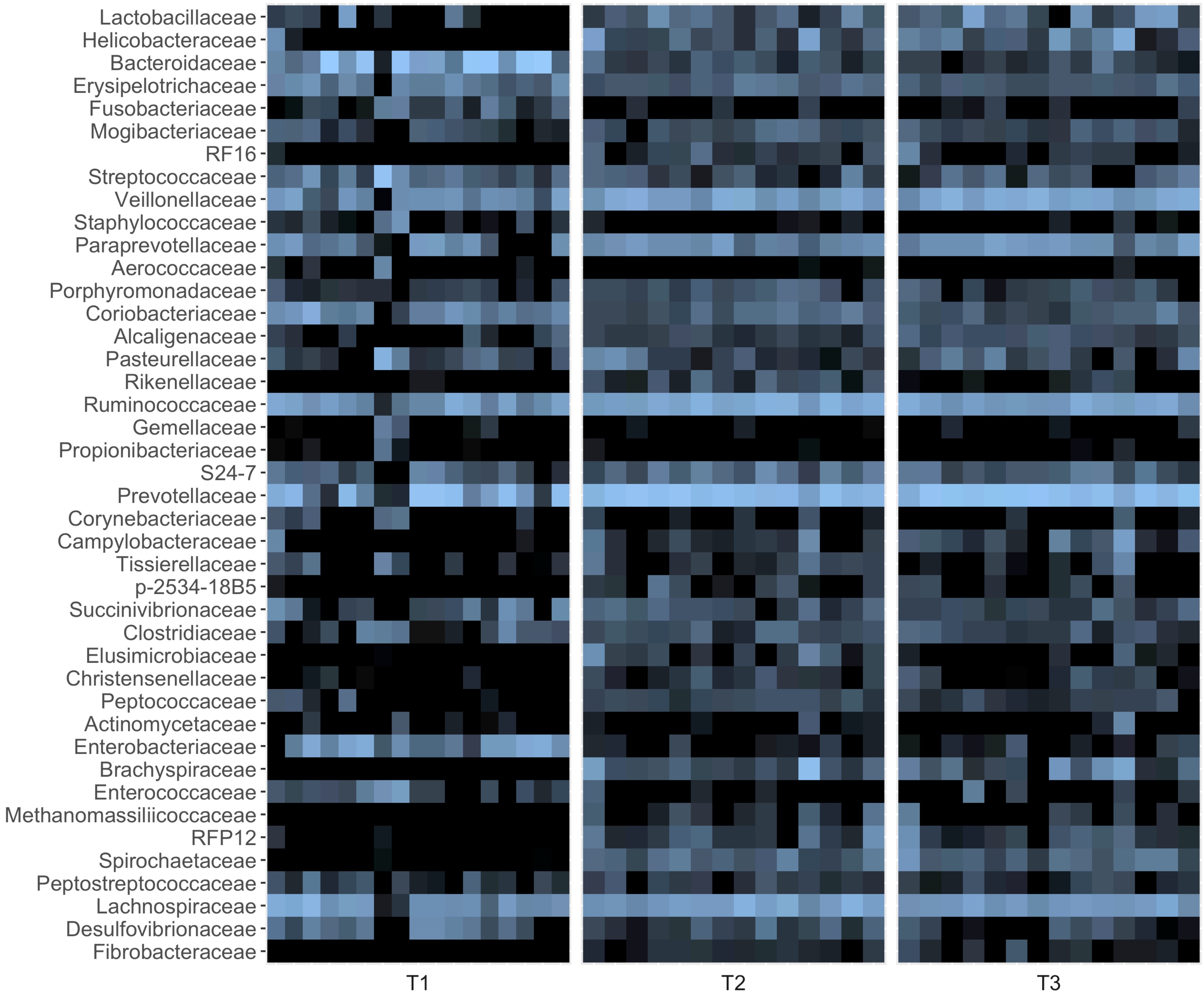

### Supplementary figure 5

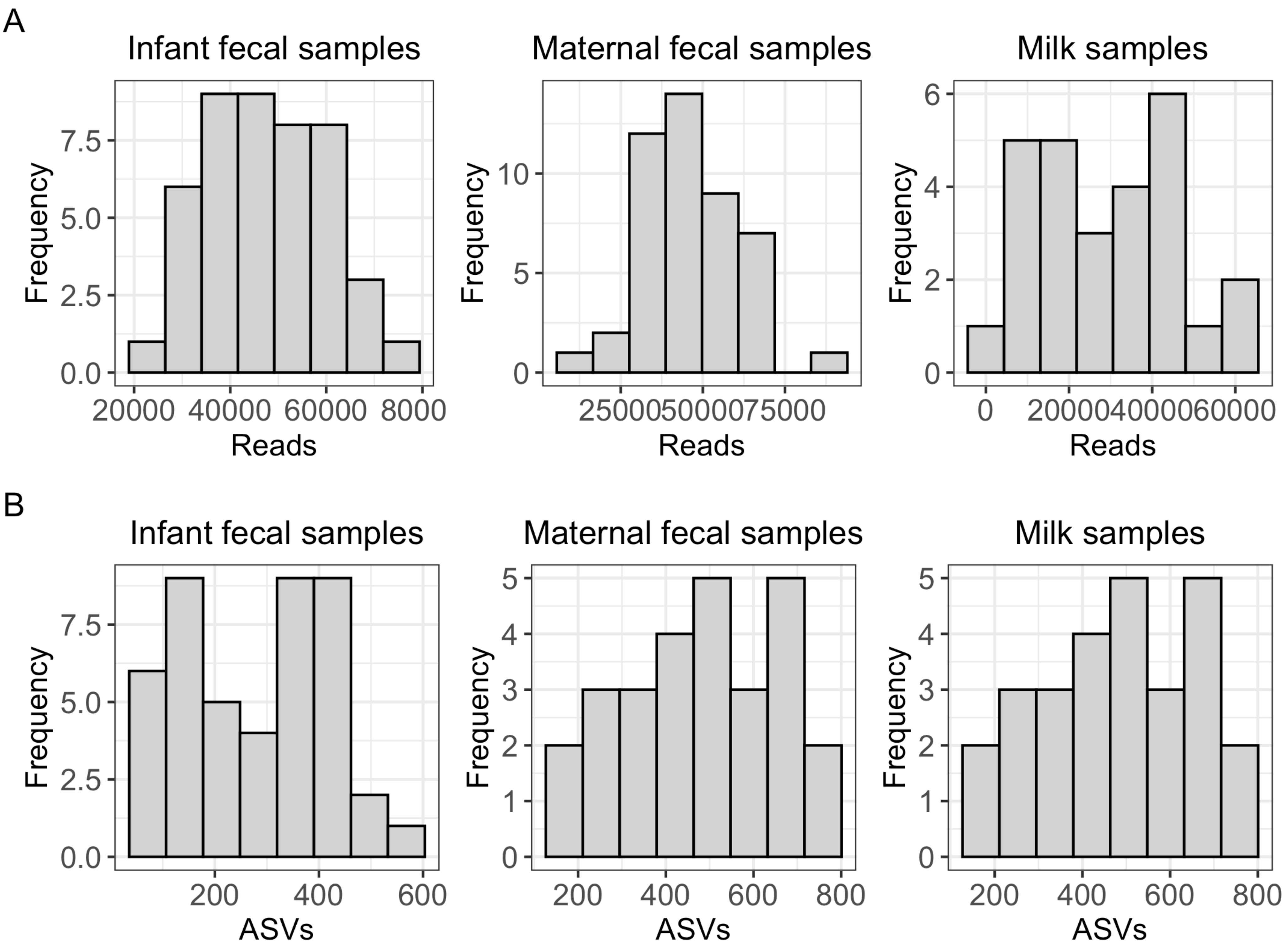
